# Mapping the thymus in the viscoelastic landscape of biological tissues

**DOI:** 10.64898/2026.03.26.714427

**Authors:** Francesco Fontana, Giulia Paties Montagner, Paolo Signorello, Arti Ahluwalia, Ludovica Cacopardo

**Affiliations:** Department of Information Engineering, University of Pisa, Largo L. Lazzarino 1, 56122 Pisa, Italy; Research Center “E. Piaggio”, University of Pisa, Largo L. Lazzarino 1, 56122 Pisa, Italy; Department of Translational Research and New Technologies in Medicine and Surgery, University of Pisa, Via P. Savi 10, 56126 Pisa, Italy; Centro 3R, Italy

**Author notes:** Corresponding author (mail).

**Keywords:** Thymus, Viscoelasticity, Microstructure, Extracellular matrix, Soft tissues

## Abstract

The thymus plays a pivotal role in the generation of immunocompetent T cells. Although its function is dependent on its complex extracellular matrix, its 3D architecture and mechanical properties remain poorly characterised This knowledge gap limits efforts to model and engineer the organ, which is a critical step towards the development of strategies for the treatment of many haematological and autoimmune diseases. Here, we provide the first comprehensive multiscale dataset of bovine thymic extracellular matrix architecture and viscoelastic behaviour, including quantitiative descriptors such as relaxation times, instantaneous and equilibrium elastic moduli, storage and loss moduli, and spatial mechanical heterogeneity. Taken together, our data define the thymus as a compliant, highly dissipative viscoelastic organ with a fibrillar architecture.

They also represent a unique database, which, for the first time, paves the way for quantitative thymus tissue engineering.

## I. Introduction

The thymus is a primary lymphoid organ and, as part of the adaptive immune system, its main function is to produce self-restricted and self-tolerant T cells [1]. It is not only dedicated to the development of T lymphocytes with specific antigen recognition receptors and long-term immunological memory, but also to the selection of those T cells which are not autoreactive (central tolerance) [2]. The thymus is divided into two main lobes, each enveloped by connective tissue (septa). Each lobe, in turn, is divided into smaller lobules, presenting two compartments: an outer one, called the cortex, and an inner one, the medulla, where early- and later-stage T cells develop, respectively [3]. These compartments are also enriched with other cell types: thymic epithelial cells (TEC), mesenchymal cells, endothelial cells, and dendritic cells. More specifically, it is up to mesenchymal cells and TEC to produce the extracellular matrix (ECM), which is mainly composed of collage type I and IV, laminin, and fibronectin, organised in an intricate 3D architecture [4], i.e. a porous and fibrillar meshwork which is critical for enabling the molecular pathways requested for T cells development [5], [6]. Despite its gradually decreasing activity with age [7], the organ is crucial for the normal development of the immune system throughout life. Since all thymic processes critically rely on the structural and mechanical features of the ECM [8], replicating its microenvironment is crucial for long-term *in-vitro* culture of thymic cells and for the development of innovative cell therapies for the treatment of immunological (e.g. thymic epithelial stem cells) and haematological disorders (e.g., CAR-T [9]).

However, most studies on the characterisation of thymic structure are only qualitative and largely based on decellularized thymic tissues employed as cell culture scaffolds. For example, Fan *et al*. (2015) [1] performed scanning electron microscopy (SEM), histology and immunostaining of the main thymic ECM proteins on sections of tissue and acellular scaffolds derived from mice. Their aim was to assess whether the decellularization process preserved the 3D fibrillar meshwork typical of thymic tissue. Asnaghi *et al*. (2021) [10] also used SEM and immunostaining to characterize the ECM microstructure from mice thymus, comparing native and decellularised tissue. The SEM images showed a highly homogeneous well-interconnected pore structure, with an unaltered ECM composition. A slightly different approach was followed by Campinoti *et al*. (2020) [11], who employed micro-computed tomography for the structural characterization of thymic samples harvested from Sprague-Dawley rats. They were mainly interested in demarcating the cortical and medullary regions through virtual segmentation, and in reconstructing a whole-organ 3D image at high resolution.

Some macrostructural parameters that are relevant from a histopathological perspective [12] have also been measured. For instance, Losada-Barragàn et al. (2019) [13], derived variations in the cortex/medulla thickness ratio (CT/MT) in thymuses harvested from mice infected with *Leishmania infantum*, to verify differences with respect to non-infected controls. Vascellari et al. (2012) [14], instead, evaluated CT/MT variations in bovine samples to with the aim of studing the influence of corticosteroids dosage (employed as growth promoters) on beef.

From a mechanical perspective, literature studies on the thymus are far fewer than for other tissues (e.g. liver, heart, bone, brain) and the organ’s viscoelastic properties have never been investigated. Some studies were performed *in vivo* through shear-wave elastography (SWE), deriving a shear wave stiffness (SWS) representative of the organ’s apparent elastic modulus. Adamczewski *et al*. (2021) [15], for example, used SWE to differentially diagnose ectopic thymuses in children, also providing quantitative values of the SWS according to different imaging planes (transverse and longitudinal). They obtained a SWS value of 7.23 ± 1.78 kPa (mean ± standard deviation (SD)) for the transverse plane; whereas, for the longitudinal plane, the relative SWS value was 11.64 ± 3.64 kPa. Bayramoğlu *et al*. (2020) [16] adopted SWE *in vivo*, on children. The SWS of thymus of all participants was 6.76 ± 1.04 kPa. However, limitations related to the use of ultrasound can affect measurements: poor acoustic window, limited penetration, and rib/lung shadows jeopardize the achievement of reliable results [17]. Asnaghi *et al*. (2021) [10], characterized thymic nanomechanical properties by means of atomic force microscopy (AFM). They derived surface force maps and visualized the spatial distribution of nanomechanical properties at micrometric resolution. The force maps of native samples revealed a recurrent pattern, possibly associated with cell compartments. The authors estimated a stiffness of 12 ± 6 kPa, attesting to the heterogenity of the samples, and perhaps also the instrumental noise associated with AFM measurements. Overall, a quantitative characterisation of native thymic tissue microstructure and viscoelastic behaviour, required to guide the design of biomaterial scaffolds and *in vitro* thymus models, is still lacking. To fill this gap, we conducted a multiscale mechanical and structural analysis of the thymus.. Mechanical characterisation included local indentation mapping at millimetric resolution together with bulk compression and shear testing. Moreover, because viscoelastic properties reported for soft tissues are strongly influenced by the mode and timescale of loading, we implemented a multimodal protocol based on step, ramp, and sinusoidal deformation to encompass a range of testing scenarios. Tissue composition was evaluated histologically at the macroscale, while the ECM microstructure, including fiber and pore size, was quantified by SEM. This work provides a first quantitative basis for defining engineering specifications for thymus models and broadens the current mechanical landscape of soft biological organs.

## II. Materials and Methods

### 1. Mechanical characterization

Viscoelastic testing of soft tissues can be performed using different methods and at different scales, and the results may differ accordingly. As the viscoelastic properties of the thymus have not been probed to date, we conducted five typologies of mechanical tests to cover a range of modalities: indentation stress-relaxation (iSR), bulk stress-relaxation (bSR), bulk epsilon-dot spectrum 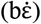, bulk sinusoidal (bSI) and bulk shear (bSH).

#### 1.1. Thymus specimen preparation

All studies were performed on bovine tissue. Despite intrinsic interspecies differences between humans and bovines, this animal was chosen not only for the similar timing of immune system development, but also because it shares many pathogens (such as coronaviruses, papillomaviruses, tubercolosis), with similarities in the immune response to them [18]. In addition, with respect to other animals intended for human consumption, bovines are more often butchered at a young age. Ten fresh thymuses from cattle (Fig. 1a.1) were kindly provided by a local slaughterhouse (Macelli di San Miniato s.r.l., Pisa, Italy). The use of the tissue did not require ethical approval, since the thymus is a slaughterhouse waste product, although the Animal Welfare Body of the University of Pisa was informed of – and endorsed - the study (Deliberation # 5 date: 16-4-2025). The samples used in this study were harvested from 12-16 months old animals, before thymic involution processes [19], which in beeves typically start at the 19^th^-21^st^ month with moderate adipose tissue infiltration in the cortex [20].

**Fig. 1.**
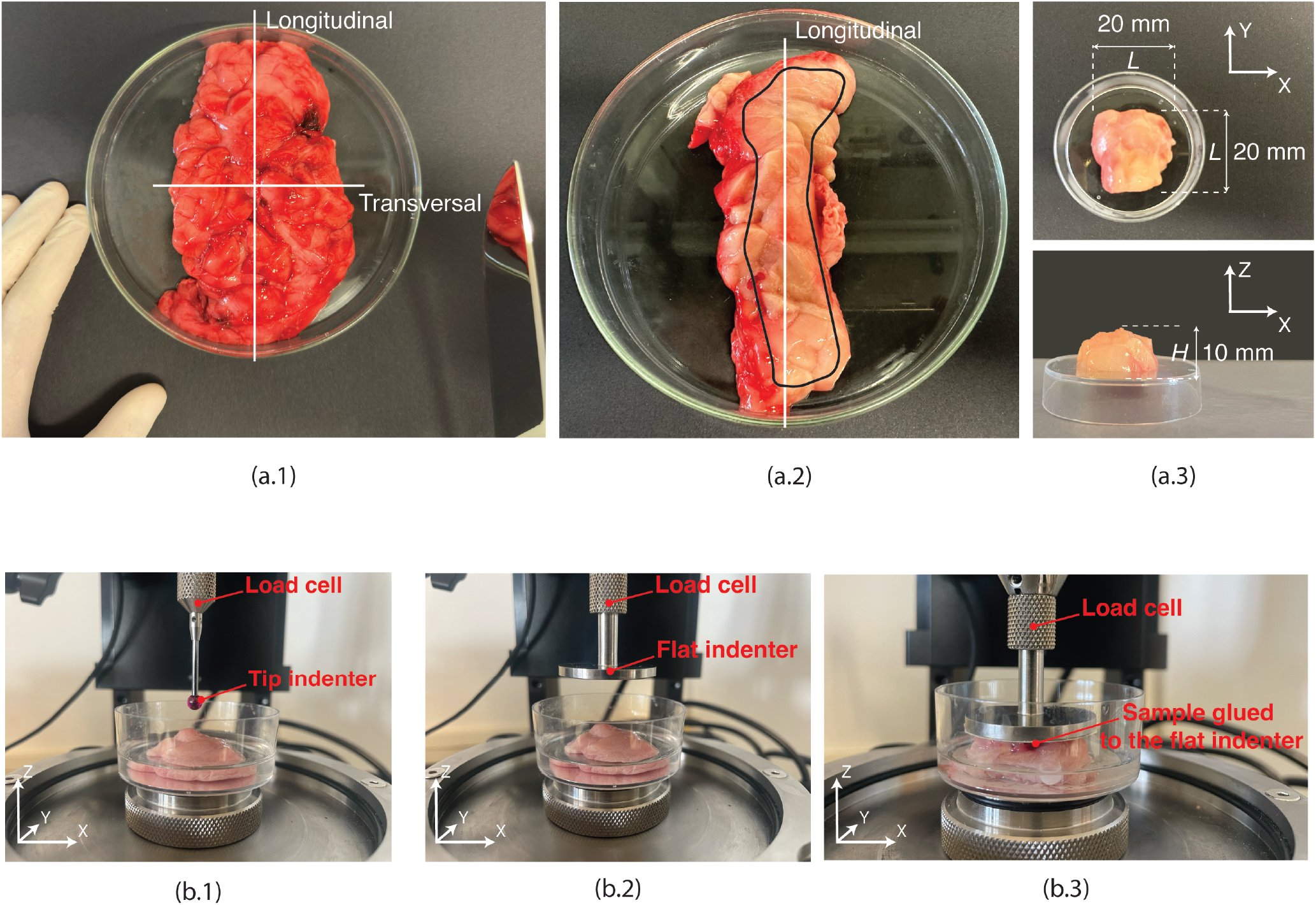
(a.1) Top view of fresh thymus, with highlighted cutting directions. (a.2) Top view of a longitudinal inner section of thymus: the black boundary defines the tissue region considered for a generic sample extraction in a longitudinal cut. (a.3) Top view (upper panel) and side view (lower panel) of the final thymic sample derived from a longitudinal inner section, with relative dimensions. (b) Testing assembly: (b.1) indentation stress-relaxation (b.2) bulk stress-relaxation/epsilon-dot spectrum/sinusoidal, and (b.3) bulk shear.

The tests were performed on fresh samples extracted from longitudinal sections (Fig. 1a.1) and only the inner region of the tissue was studied to avoid any superficial and connective portion (Fig. 1a.2). Six to seven samples (base area *L*^*2*^ ∼ 20 ⨯ 20 mm; height *H* ∼ 10 mm, Fig. 1a.3) were manually excised with a scalpel from each organ. At least three samples from three different organs were used for each mechanical test.

Tools and gloves were kept humidified throughout the preparation and mounting phase, to avoid any adhesion and tearing while handling. Nevertheless, due to the particularly soft nature and lobular architecture of thymic tissue, specimens tended to deform under their own weight when manipulating and moving them, which led to a variation in final sample dimensions with a tolerance of ± 3 mm for both base edges and heights. Engineering stresses and strains were thus calculated considering the dimensions of the samples measured after positioning them in the testing rig. Further experimental details on tissue handling, storage, size measurement and testing boundary conditions are reported in SI1. The influence of tissue freezing-thawing and tissue excizing direction (longitudinal and transversal) are shown in SI2 and SI3, respectively.

#### 1.2. Experimental set-up and mechanical tests

After extraction, each sample was fixed with cyanoacrylate glue at the center of a plastic petri dish and half-immersed in PBS (phosphate buffered saline) to keep it hydrated during the entire test procedure. Glueing the sample bottom surface did not impose overconstraining conditions during compression tests, as the upper and lateral sample faces were able to deform. The petri dish itself was secured to a metal sample holder with adhesive tape. This assembly was finally mounted on the mechanical tester, as represented in Fig. 1.b. All mechanical tests were performed using a Mach-1 v500css (Biomomentum Inc., Montreal, Canada) equipped with a 17 N multiaxial load cell. The parameters adopted for each of the tests performed are listed in Table 1 and were taken from reference values for soft tissues, in timescales relevant for organ engineering. Data from poroelastic investigations were not considered since the loading times employed here are much lower than the characteristic timescale for soft tissue poroelasticity [21], [22], [23]. For indentation and bulk tests (Fig. 1b.1 and Fig. 1b.2, respectively), the contact finding was achieved by setting a contact force threshold of 0.04 N; then, data were acquired with the tip/flat indenter starting at a precaution distance of 5 mm from the tissue sample surface, to guarantee a zero pre-stress initial condition and a constant velocity before contact. For the indentation tests, the sample surface orientation at an xy position was automatically detected by the Mach-1 through the least squares method, allowing identification of the normal vector and indenting along that direction. In the case of shear tests, the sample was glued to the flat indenter with a contact threshold force of 0.04 N, to avoid sample slipping during the experiment, and the recording of friction forces rather than shear-generated ones (Fig. 1b.3) [24]. In the case of step and ramp inputs, a generalized Maxwell (GM) model was used to derive the viscoelastic descriptors: instantaneous and equilibrium elastic moduli (E_inst_ and E_eq_), as well as the characteristic relaxation time(s) (τ). For local indentation tests, E_inst_, E_eq_ and τ distribution maps were also extrapolated for each sample. For sinusoidal stimuli, the storage and loss moduli (E’ and E”), as well as the phase shift angle (δ) were calculated. Finally, the shear apparent elastic moduli (G_app_) was obtained.

**Table 1.**
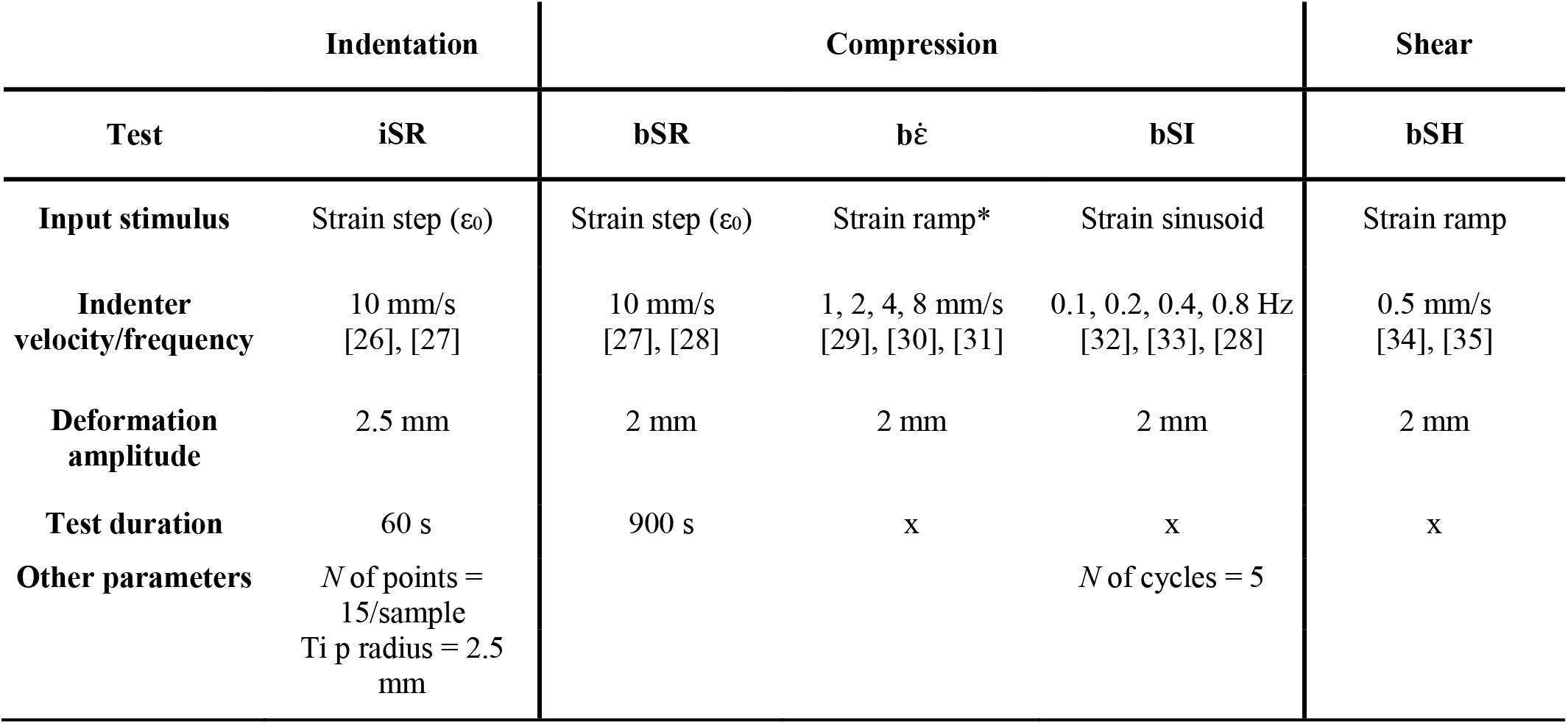
Parameters used for the different testing modalities. *N* = number; iSR = indentation stress-relaxation; bSR = bulk stress-relaxation; 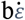 = bulk epsilon-dot spectrum; bSI = bulk sinusoidal; bSH = bulk shear. *based on a 5% LVR.

#### 1.3. Data analysis and constitutive modeling

For bSR, 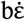, bSI and bSH tests, force and deformation data were normalized with respect to sample cross-sectional area (*L*^*2*^) and sample height (*H*), respectively, to obtain experimental engineering stress and strain (σ, ε) (Fig. 2a,b,d)). In the case of b and bSH tests, the apparent elastic and shear moduli (E_app_ and G_app_, respectively) were derived as the slope of the σ-ε curves in the linear viscoelastic region (LVR), i.e. the region in which stress varies linearly with applied strain (R^2^ > 0.99). The iSR data were processed considering a Hertzian contact (Fig. 2c) [25].

**Fig. 2.**
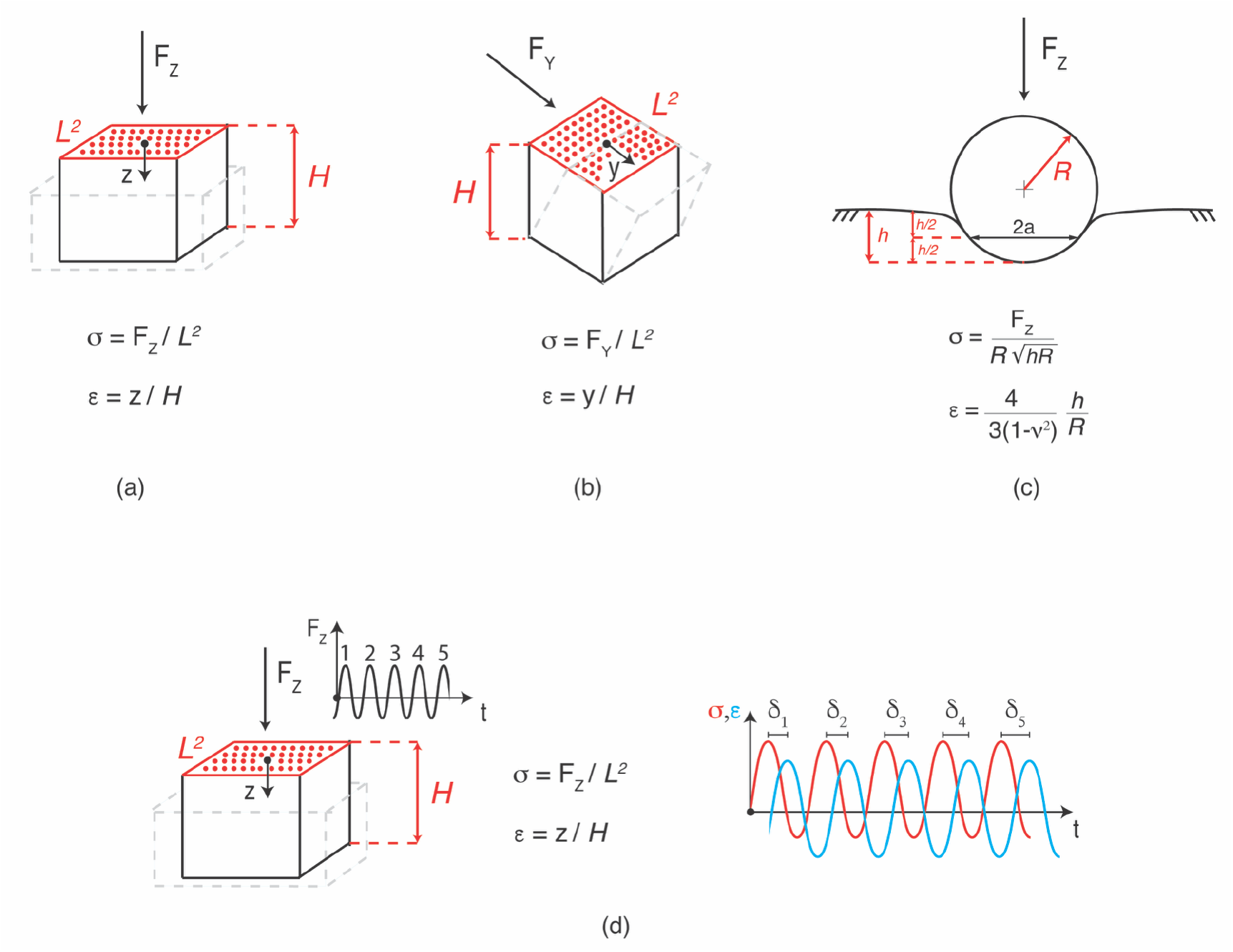
Schematic representation of test modes: (a) bulk stress-relaxation (bSR)/epsilon-dot spectrum 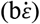, (b) bulk shear (bSH), (c) indentation stress-relaxation (iSR) and (d) bulk sinusoidal (bSI). σ = stress; ε = strain; δ = phase shift. Bulk tests are performed with a flat indenter, while local tests with a spherical tip indenter.

Standard Linear Solid models are analytically tractable and widely used for representing soft tissues, particularly at low strains. A second order (*n* = 2) GM model (Fig. 3a) – which ensures goodness of fit, while avoiding overparameterization - was thus used to represent the viscoelastic behaviour of thymic tissue across the different testing modalities (an example of how R^2^ increases with model order is shown in SI4). For iSR, bSR and b, the characteristic equations (summarized in Table 2) were fitted to the experimental data to derive the model parameters (i.e., E_0_, E_1_, E_2_, η_1_, η_2_ as depicted in Fig. 3a) using Matlab_R2023B-Mathworks. The viscoelastic descriptors E_inst_, E_eq_, and τ_i_ were calculated as reported in Fig. 3b. For b/iSR only the longer relaxation time (herein referred to as τ) was considered as it is more indicative of bulk tissue relaxation (τ1 and τ2 are reported in SI5). Differently, in the case 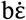, only τ_1_ was extrapolated (herein referred to as τ), as *n* = 2 caused overfitting. Surface distributions of E_inst_, E_eq_, and τ from the iSR tests were also represented by means of colormaps through the “Mach-1 Analysis” software.

**Table 2.**
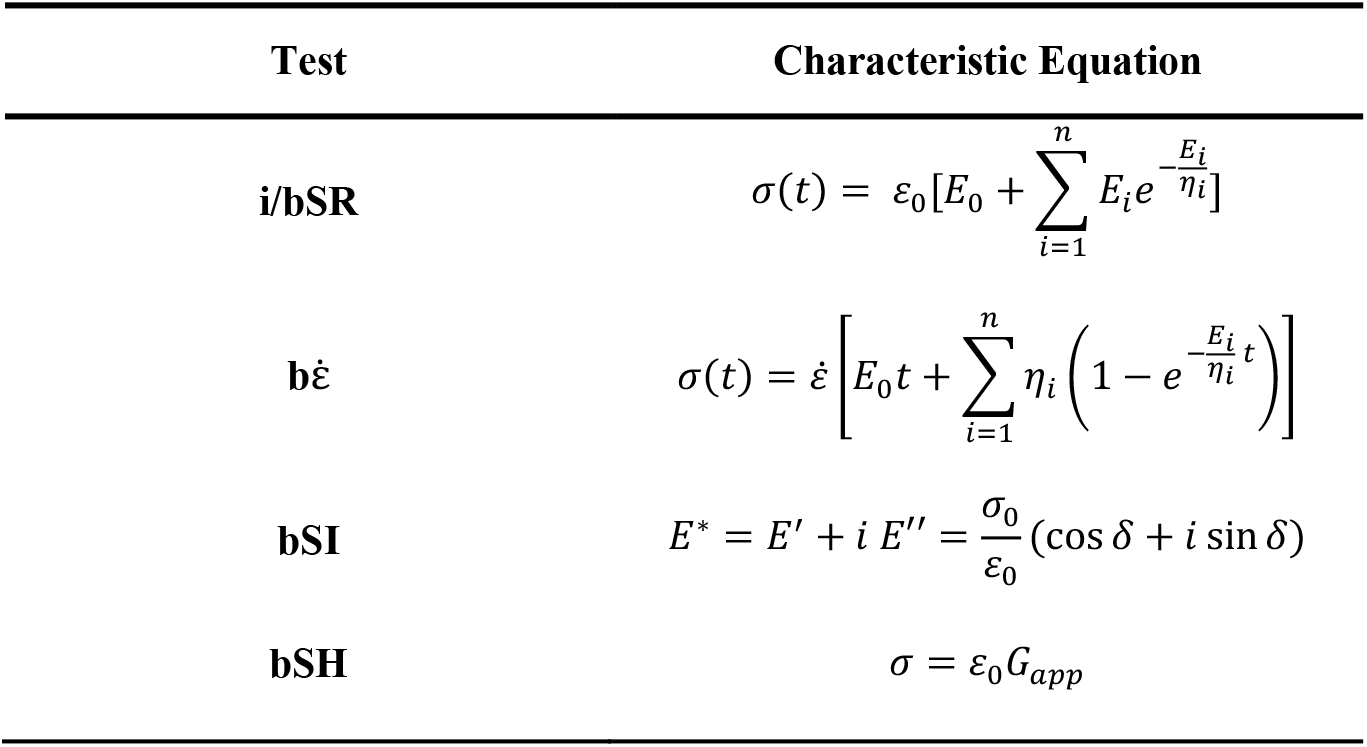
Models and equations for data analysis [36], for each mechanical test. i/bSR = indentation/bulk stress-relaxation; 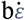 = bulk epsilon-dot spectrum; bSI = bulk sinusoidal; bSH = bulk shear.

**Table 3.** Summary of thymus multimodal and multiscale mechanical characterization expressed as median (25, 75%). All datarefer to fitting obtained with R^2^ > 0.99.

| | Indentation stress-relaxation (iSR) | Bulk stress-relaxation (bSR) | Bulk epsilon-dot (b$\dot{\epsilon}$) | Bulk sinusoidal (bSI) 0.1-0.8 Hz |
| --- | --- | --- | --- | --- |
| $E_{\text{inst}}$ [kPa] | 11.4 (8.3, 15.6) | 3.9 (2, 5.2) | 4.8 (4.2, 5.3) | x |
| $E_{\text{eq}}$ [kPa] | 2.2 (1.5, 3.3) | 0.8 (0.4, 1.2) | 1.4 (1.3, 1.7) | x |
| $\tau$ [s] | 11.2 (9.7, 12) | 215.5 (177.6, 287) | 0.08 (0.06, 0.08) | x |
| $E'$ [kPa] | x | x | x | 3.3 (3.2, 3.4) |
| $E''$ [kPa] | x | x | x | 1.1 (0.8, 1.2) |

**Table 4.** Structural and histological quantities derived from SEM and optical microscopy image analyses.

| | MT [mm] | CT [mm] | PT% [%] | AT% [%] | CT/MT [-] | CA/MA [-] | PD [μm] | FD [μm] |
| --- | --- | --- | --- | --- | --- | --- | --- | --- |
| **Mean**<br>**± SD** | 2.1 ± 0.7 | 3.3 ± 1.3 | 83 ± 4 | 7 ± 3 | 1.6 ± 0.5 | 7 ± 1 | 1.2 ± 0.4 | 0.2 ± 0.1 |

**Fig. 3.**
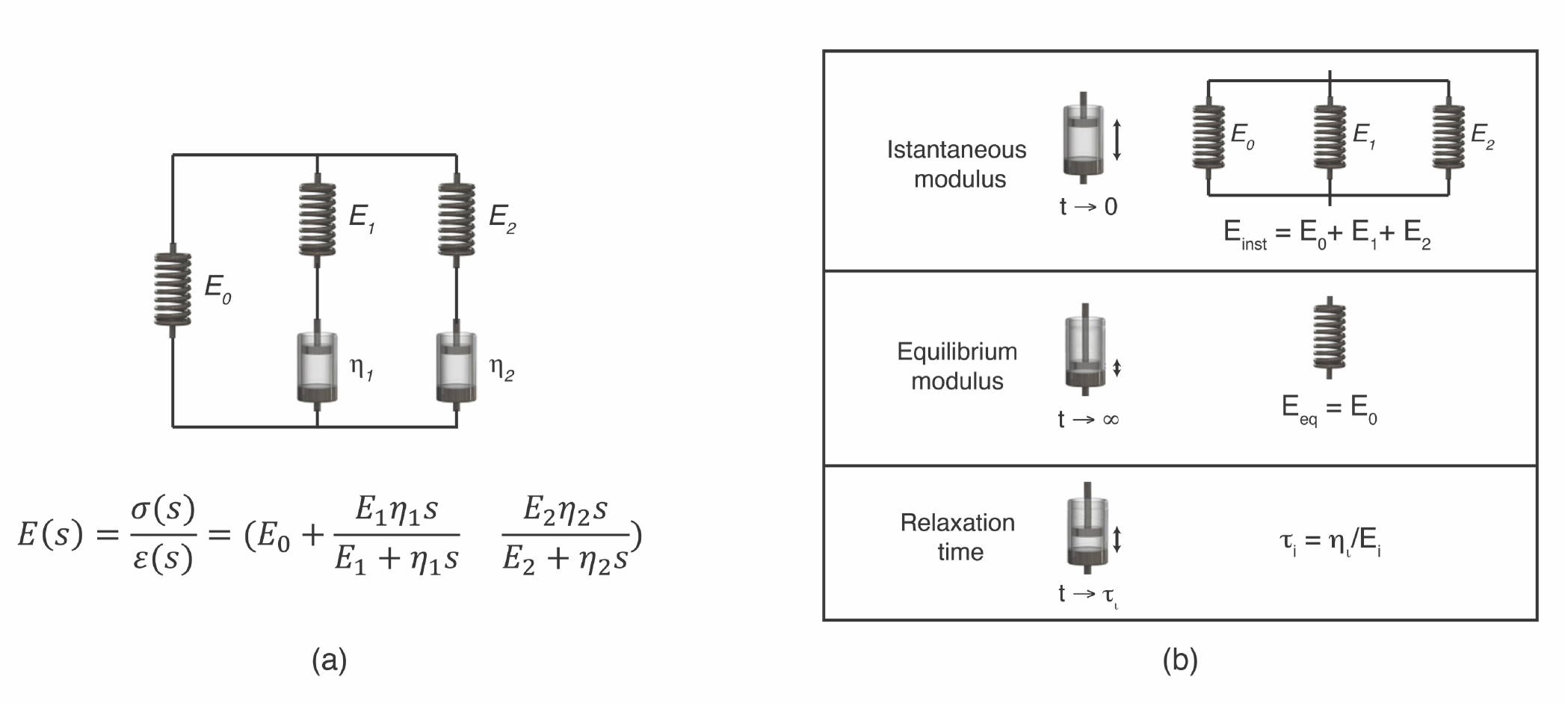
(a) Generalized Maxwell model for *n* = 2, and its transfer function in the Laplace domain for solid materials; (b) Viscoelastic descriptors as a function of the Generalized Maxwell model parameters (for *n* = 2). Adapted from [36]

In the case of sinusoidal stimuli in bSI tests, the phase shift (δ) was calculated as illustrated in Fig. 2d and the complex elastic modulus (E^*^) was derived as reported in Table 2, defining the storage and loss moduli E’ and E” respectively as the real and imaginary part of E^*^.

### 2. Structural characterization

As regards the structural analysis, at a macroscale, we analysed thymic tissue composition by quantifying the following parameters: the lobular percentage of adipose tissue infiltrations (AT%) and parenchymal areas (PT%) (i.e., the lobular percentage of cortex and medulla), as well as the cortex/medulla area ratio (CA/MA), wich was defined as the ratio between total cortex and medulla areas, in all analyzed lobules. Furthermore, the cortex and medulla mean thicknesses (CT and MT, respectively) were derived, evaluating the CT/MT. Finally, at the ECM microscale, we derived the diameter of fibres and pores (FD and PD, respectively), which had never been investigated in past literature.

#### 2.1. Scanning electron microscopy and image analysis

Fresh samples were fixed with 10% w/w formalin solution for 3 days, then washed with d-H_2_O overnight, at room temperature. After dehydration in graded ethanol solutions (from 70% to 100% EtOH), they were embedded in paraffin. Cubes of 2 mm in sides were deparaffinated, rehydrated (from 95% to 100% d-H_2_O), freeze-dried and finally sputter-coated with platinum before examination with a FEI ESEM Quanta 450 FEG microscope. Thymic micro-morphology was observed qualitatively and PD and FD assessed quantitatively using ImageJ [37].

#### 2.2. Histology and image analysis

Deparaffinized and rehydrated sections of 5 μm thickness were stained with hematoxylin & eosin to visualize nuclear content and tissue microstructure. The CT, MT and CT/MT, as well the AT%, PT% and CA/MA, were manually quantified by using ImageJ. More specifically, in the case of CT and MT, after selecting a lobule, the maximal width of the region of interest was manually drawn with a line graduated on the image scale bar (starting and ending at the interlobular connective or adipose tissue for CT, and at the surrounding cortex for MT) [14]. In the case of AT%, PT% and CA/MA, images were binarized in a segmentation step, with grey levels ranging from 0 (white) to 1 (black). Each type of tissue was associated with a specific grey shade and quantified through thresholding [38], [39].

### 3. Statistical analysis

Data from mechanical and structural characterization underwent a Kolmogorov-Smirnov normality test with significance threshold set at *p* = 0.05. Non-normal distributions are reported as boxplots showing the median value and the 25^th^ and 75^th^ quartiles, with whiskers ranging from the minimum to the maximum value. Normal distributions are reported as bar plots. When comparing two non-normal distributions, Mann-Whitney tests were performed to identify statistically significant differences. For more than two non-normal distributions, Kruskal-Wallis with Dunn’s post-hoc test analyses were performed. The significance threshold was set at *p* = 0.05. All analyses were performed using GraphPad Prism 10.

For the analysis of SEM and histology images, at least 3 regions of interest were captured for each tissue sample (*N* of tissue samples = 3). In the case of SEM images, for each region at least 20 pores and fibres were analysed. For histology images at least 10 lobules were analysed.

## III. Results and Discussion

There is growing interest in developing biomaterial-based scaffolds that can recapitulate the thymic microenvironment, with greater reproducibility, tunability, and translational potential. To date, however, most efforts to engineer thymic tissue have relied on decellularized extracellular matrices derived from animal or donor organs. While they provide biologically relevant cues, they are limited by availability, batch variability, and the possibilty of ECM damage or cytotoxic residues arising from the decellularization process. The rational design of fit for purpose and reliable biomaterials requires a quantitative understanding of the native thymus as a mechanical and structural template, particularly with regard to its viscoelastic behaviour. The multiscale and multimodal characterization presented here helps fill the gap by defining a set of mechanical and structural specifications for engineering the thymus.

We first assessed whether sample preservation and tissue orientation affected viscoelastic descriptors, comparing fresh and thawed samples and longitudinal and transversal sections. In the case of iSR tests, significant differences were observed between fresh and thawed thymus (see SI2), whereas no significant differences were observed between longitudinal and transversal sections (see SI3). All tests were thus conducted on fresh samples. For the iSR tests, maps and viscoelastic descriptors distributions are reported in Fig. 4. Although boxplots derived from the overall samples showed a small interquartile range for each viscoelastic descriptor (Fig. 4a), the surfaces were essentially heterogenous, as reflected in the histological images (Fig. *6*). In fact, as the samples were interrogated with small tips (radius = 2.5 mm), we observed a high degree of intratest variability: in each sample the values of viscoelastic descriptors ranged over an order of magnitude along the sample surface (e.g., E_inst_ and E_eq_ in the representative sample, Fig. 4b).

**Fig. 4.**
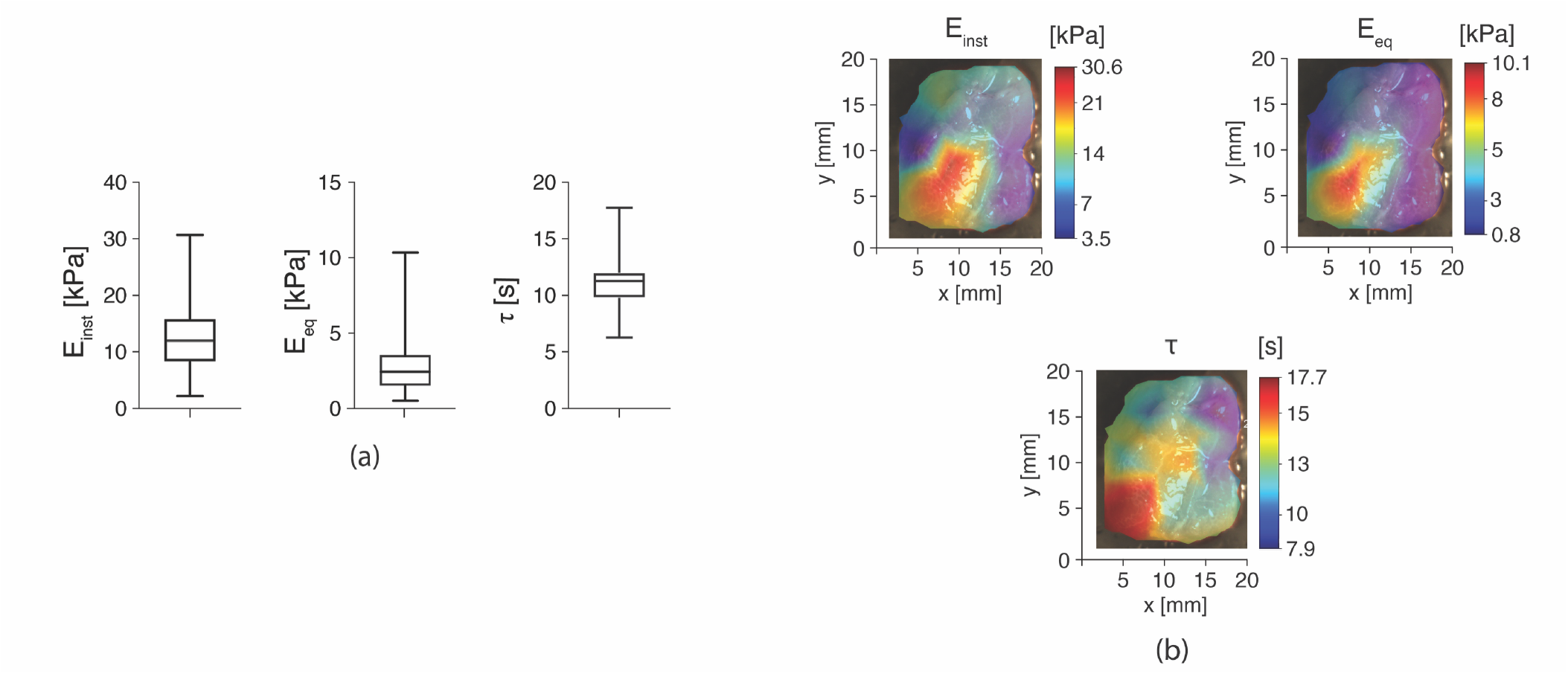
(a) Instantaneous elastic modulus (E_inst_), equilibrium elastic modulus (E_eq_) and characteristic relaxation time (τ), for all samples tested with iSR. (*N* of samples = 4; *N* of points/sample = 15) (b) Surface plots of E_inst_, E_eq_, and τ for a representative sample.

For the bSR and 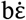 tests (Fig. 5.a1 and Fig. 5.a2), the estimated values of E_inst_ and E_eq_ were statistically comparable (E_inst_ =3.9 kPa for bSR *vs* E_inst_ =4.8 kPa for 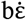 (medians), *p* = 0.3; E_eq_ =0.8 kPa for bSR vs E_eq_ =1.4 kPa for 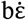 (medians), *p* = 0.06). Nevertheless, significant differences in the characteristic relaxation times were observed for the two testing modalities (τ = 215.5 s for bSR *vs* τ = 0.08 s for 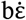 (medians), *p* = 0.008). That τ varies for different testing modalities has been reported previously and can likely be ascribed to intrinsic differences in the input stimuli, which probe different time windows, fluid phase redistribution and structural rearrangements.

**Fig. 5.**
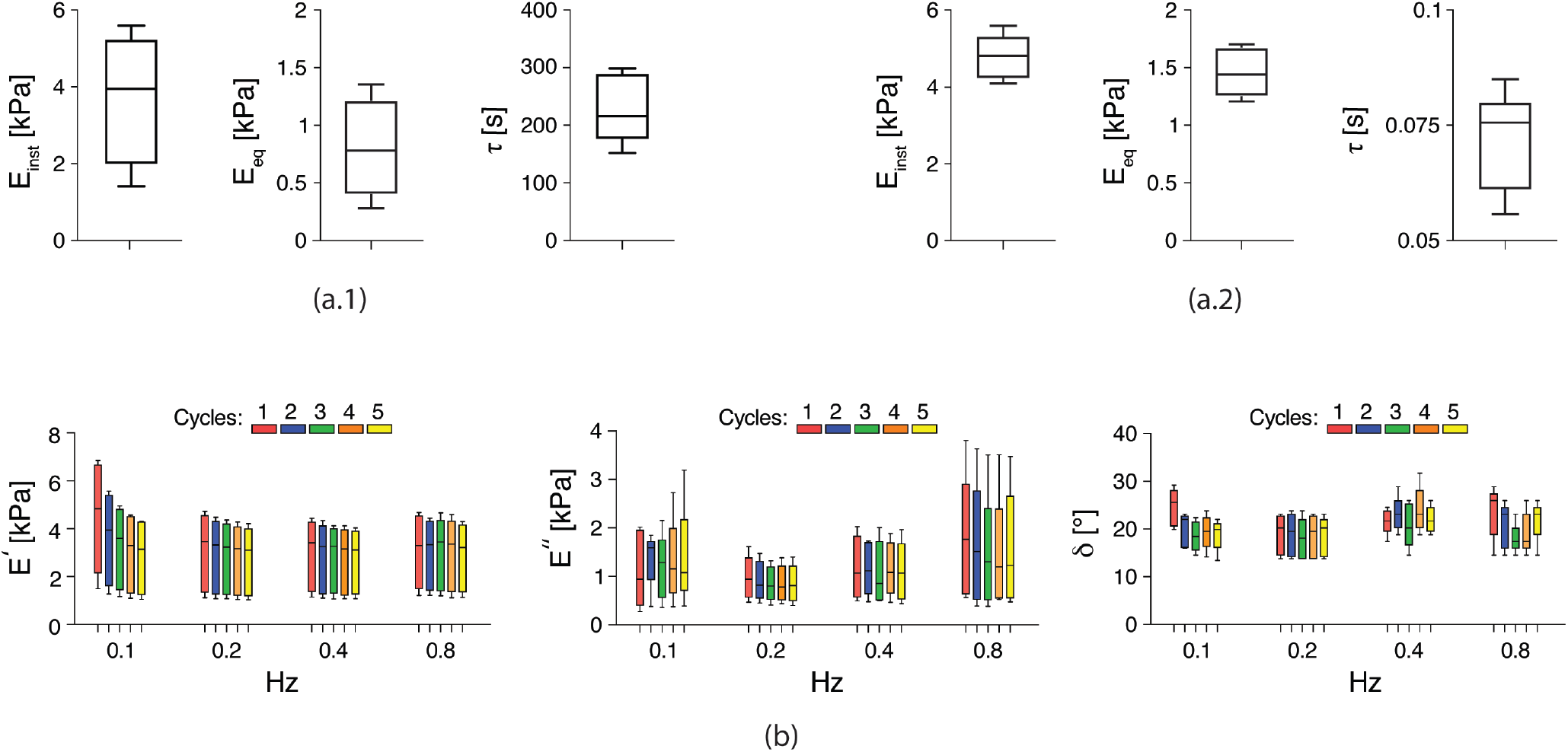
Instantaneous elastic modulus (E_inst_), equilibrium elastic modulus (E_eq_), and characteristic relaxation time (τ), for all samples tested with (a.1) bSR and (a.2) 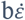 (*N* of samples = 4). (b) Storage modulus (E’), loss modulus (E’’), and phase shift (δ) for each cycle (*N* of cycles = 5) and tested frequency (0.1, 0.2, 0.4, 0.8 Hz) (*N* of samples/frequency = 4). Kruskal-Wallis followed by Dunn’s post-hoc test showed no statistically significant differences between cycles or frequencies tested (GraphPad Prism 10).

This protocol dependence is further compounded by practical limitations: 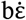 relies on a finite set of strain rates and therefore samples only a limited portion of the spectrum of relaxation processes (this also justifies the overfitting for *n* = 2 in the GM), whereas bSR experiments cannot realize an ideal step strain and are necessarily performed over a finite observation time [40]. Relaxation times (i.e., τ_1_ and τ_2_) for bSR and 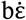 tests are reported in SI5.

When comparing results between iSR and bSR, statistical differences were observed in terms of both E_inst_ (11.4 kPa for iSR vs 3.9 kPa for bSR (medians), *p* = 0.0005) and E_eq_ (2.3 kPa for iSR vs 0.8 kPa for bSR (medians), *p* = 0.003). Variations between bulk and local compression tests are well-documented, particularly for biological tissues. This discrepancy is generally attributed to factors related to geometry, boundary conditions and strain field heterogeneity differences [41], [42], [43],[44], [45]. Surface roughness and sample heterogeneity may also affect local mechanical properties given the differing fractions and distributions of mechanically influential components such as collagen (type I and type IV), fibronectin and fat deposits.

On the other hand, variability in 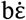 (Fig. 5,a2) is minimal, probably due to the global fitting of a higher number of testing curves from different strain rates with respect to bSR (Fig. 5,a1). Moreover, when applying slower strain rates in the 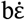 method, the initial transient is smoother and less sensitive to precision control and inertial artifacts [46], [47]. The variability in bSR tests might be due to the strong sensitivity of tissues to ramp speed. This effect has been previously observed: the faster the ramp, the more sensitive the mechanical response becomes to small differences in ramp duration, actuator inertia and precision control [48], [49]. Moreover, at high deformation rates, sample response is strongly affected by fluid pressurization and fiber recruitment [50], [51], which, considering the instrinsic heterogeneity of the thymic tissue, may lead to higher inter-sample variability. Finally, rapid compression produces sharp transients that may not be well captured by lumped parameter model descriptions and extracted parameters may thus be more sensitive to noise [52].

It is also worth mentioning that, although the E_inst_ values from iSR, bSR and 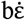 tests reported here are quantitatively comparable to SWS values previously mentioned in the state of the art (Adamczewski *et al*. (2021) [15] and Bayramoğlu *et al*. (2020) [16]), a direct comparison between them might be misleading as they describe different mechanical behaviour under different assumptions. In fact, the two studies are essentially different to the viscoelastic tests performed in this work in terms of timescales, strain rates and frequencies. Similarly, albeit the thymic stiffness values from AFM measurements in Asnaghi *et al*. (2021) [10], are comparable to the E_inst_ values from our iSR tests, the similarity is casual.

The results for E’, E’’ and δ are reported in Fig. 5b for each cycle and frequency tested. No statistically significant differences emerged when comparing multiple frequencies and cycles for each viscoelastic parameter. A slight, but non-significant decrease in E’ with increasing number of cycles, was observed at each frequency and is likely associated with issue preconditioning effects [53]. Reference results in the investigated frequency and cycle range were: 3.3 (3.2, 3.4) kPa for E’, 1.1 (0.8, 1.2) kPa for E’’ and 20.2 (19.4, 23) ° for δ (median (interquartile range)).

Finally, G_app_ from bSH was evaluated: 4.4 kPa (4.4, 4.5) (median (interquartile range)). However, comparing bulk compression and shear tests performed at the same strain rate (0.5 mm/s) is challenging because of different experimental set-up arrangements (described in section 1.2) which are probably associated with different involvement of ECM components and consequently result in different trends, as shown in SI6.

Representative experimental curves for each test are also reported in SI7.

SEM images of thymic microstructure are reported in Fig. 6a, also showing the detection and distribution of the ECM PD and FD. Representative histologic images of thymic compartments are reported in Fig. 6b, with zoomed examples of detected and measured quantities. The SEM images confirmed the fibrous organization already observed by Fan *et al*. for mice [1]. In our work, pore and fiber diameter distributions were also calculated, which have not been previously considered for characterizing thymic tissue *ex vivo*. Regarding the histological analyses, on examining cattle beeves, Sebastianelli *et al*. [20] observed that thymus atrophy can be related to animal age. Our results are consistent with their findings, both in terms of CT/MT ratio and adipose tissue infiltration. In particular, our CT/MT ratio was 1.6 ± 0.5 (mean ± SD), which is in accordance with the median calculated in ref [20] for animals with same age as in this study. Our adipose tissue infiltration data (7 ± 3 %, mean ± SD) are also in line with the thymus atrophy score identified for this age. Furthermore, we augmented the structural characterization dataset by defining and calculating an additional parameter, CA/MA, derived from cortex and medullary areas of lobule compartments, and evaluating PT% and AT%. These additional characteristics should allow closer replication of the ECM.

**Fig. 6.**
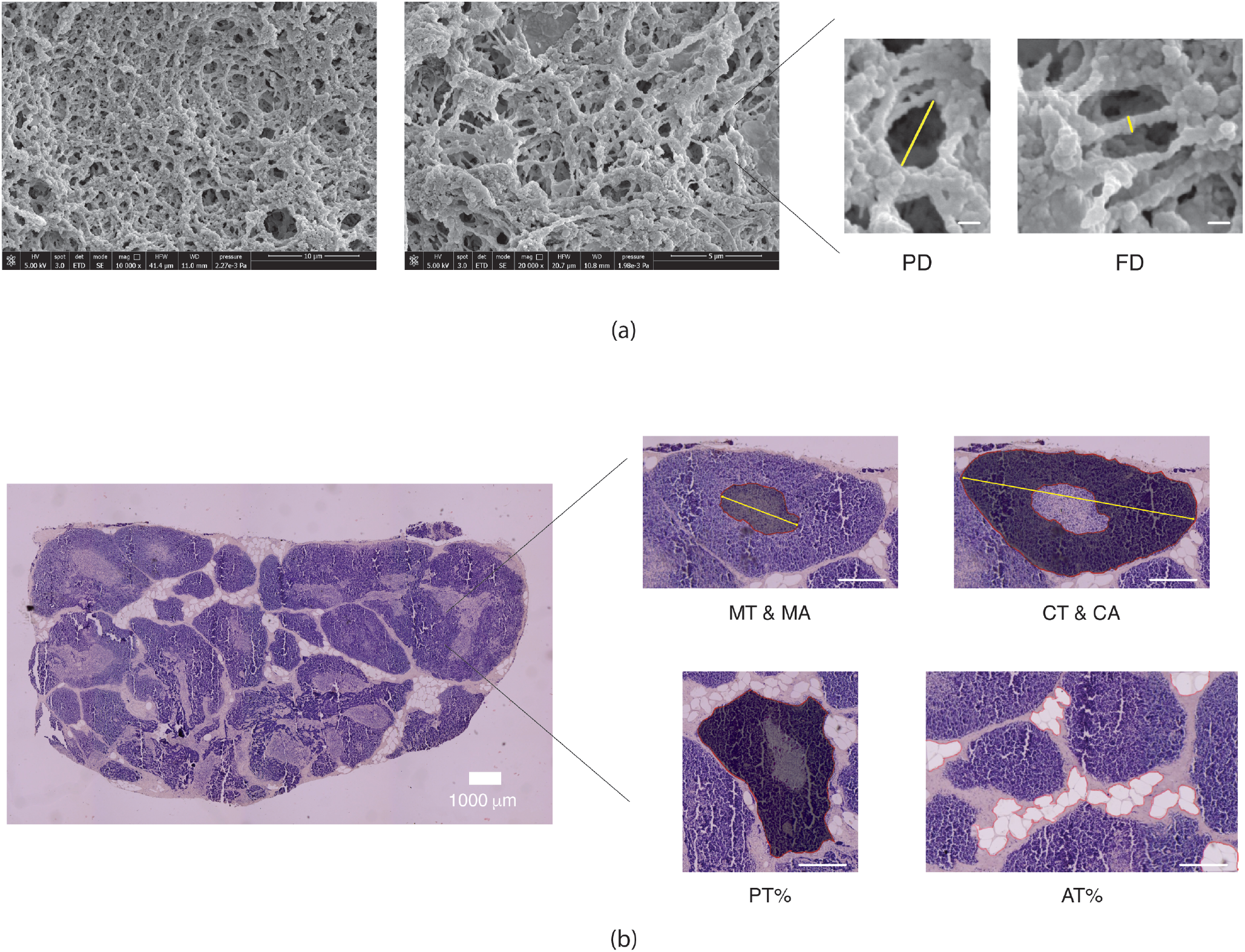
(a) Scanning electron microscopy images of thymic sections, with zoomed pore and fibre diameters (PD and FD, respectively, highlighted in yellow, white scalebar 0.1 μm). (b) Representative image of haematoxylin & eosin-stained sections of thymic lobes, with zoomed examples of detected parameters: medulla thickness (MT, yellow line) and medulla area (MA, grey-scaled area); cortex thickness (CT, yellow line) and cortex area (CA, grey-scaled area); parenchymal tissue percentage (PT%), defined as the percentage ratio between grey-scaled area (MA & CA) and total area; adipose tissue percentage (AT%), defined as the percentage ratio between grey-scaled area (adipose cells) and total area (white scalebars 0.5 mm). The CT/MT ratio was defined as the ratio between CT and MT and the CA/MA ratio was defined as the ratio between the total CA and MA.

For ease of use in biomaterial design, Table *3* and Table *4* summarise the principal mechanical and structural descriptors identified here and translate them into an initial set of design specifications for engineering thymus-mimetic scaffolds and in vitro models.

## IV. Conclusions

We provide the first multiscale and multimodal mechanical characterization of the thymus, deriving a set of viscoelastic descriptors (instantaneous and equilibrium elastic moduli, storage and loss moduli, and characteristic relaxation times). Considering different loading protocol and timescales provides researchers with a comprehensive overview of the different viscoelastic phenomena in the tissue. Beyond filling a major knowledge gap, this work delivers an initial quantitative specification set for the engineering of thymus-mimetic biomaterials and in vitro models.

Using this dataset, thymus-mimetic scaffolds can be designed and manufactured for culturing patient thymic cells, paving the way for patient-relevant preclinical testing of thymic regenerative strategies and treatments for immune and hematological dysfunctions.

## Acknowledgements

We thank Massimo Ciardelli for the technical support in the histological procedures, Irene Valente for collaborating in the mechanical experiments and Nicole Guazzelli for her precious advice for image analysis.

## Ethical statement

The authors have nothing do declare.

