## Supplementary material for "Mapping the thymus in the viscoelastic landscape of biological tissues"

### Annex 1 – Supplementary information

Received xxxxxx
Accepted for publication xxxxxx
Published xxxxxx

1

 © xxxx Nature Portfolio

#### SI1. Sample preparation and testing details

After retrieving thymic organs from the local slaughterhouse, they were brought to the laboratory in a coolbox and immediately cut (on the same day), according to the procedures described in the main manuscript. After cutting, samples were stored in the fridge, at 4°C, in a petri dish sealed with parafilm, for a maximum of five days, during which the tests were carried out. After this period they were discarded and replaced with newly procured tissue.

Sample dimensions were measured after placement in the testing rig, averaging over 6 different points on each sample. In all cases the strain is the engineering strain, while the stress is the engineering stress, with reference to these initial (average) dimensions. The intra-sample variability in dimensions was 7% maximum.

*Testing boundary conditions*

In the case of compressive stress tests (bSR, bSI and b$\dot{\varepsilon}$), only the bottom surface of the tissue was glued and fixed, whereas the upper and lateral faces were able to deform under compression, preventing overconstraining conditions through changes in specimen height and length, as suggested by Budday et al. (2017) [1].

As regards shear loading, we aimed at achieving a "no compression" condition as closely as possible through the following protocol: the flat indenter contact surface was coated with adhesive cyanoacrylate glue; then, the sample top surface height was detected by setting a contact force threshold of 0.04 N. In this way, contact detection and sample sticking were achieved at the same step. Following this step, the shear test was performed, respecting well-established boundary conditions as reported in the literature [1], [2], [3].

Specifically:

- the bottom surface of the specimen was glued and fixed to prevent upward or horizontal movement;
- the top surface, glued to the flat indenter, was controlled in its horizontal movement through the flat indenter itself, recording the returned force by means of the load cell;
- The glued top surface, fixed in height through the flat indenter, guaranteed a constant vertical distance.

As regards Hertzian contact, the "Normal Indentation" function of the "Mach-1 v500css" mechanical tester, was adopted. This allows the instrument to detect the contact and orientation of the surface at an XY position using the Least Squares Method to find the best-fitting plane to a series of 5 points with (x,y,z) coordinates; then, it finds the normal vector and indents along that direction, starting at a precaution distance of 5 mm from the tissue sample surface, to guarantee a zero pre-stress initial condition and a constant velocity before contact.

#### SI2. Freezing-thawing influence

After samples collection from longitudinal cutting sections, some of them were kept at -23 °C, for at least 24 h. Then, they were thawed at 30 °C and were mechanically tested with indentation and bulk stress-relaxation. Fig. S1, Fig. S*2* report data comparisons with fresh samples collected from the same cutting direction (longitudinal), applying Mann-Whitney statistical test (GraphPad Prism 10 - *N* of samples per test = 4; *N* of points per surface, for indentation = 15*).* Althought bulk tests show no significant differences in the viscoelastic descriptors, significant differences can be observed for both $E_{inst}$ and $E_{eq}$ in the case of indentation stress-relaxation tests. At the local scale, this can be ascribed to local tissue alterations due to freezing/thawing process.

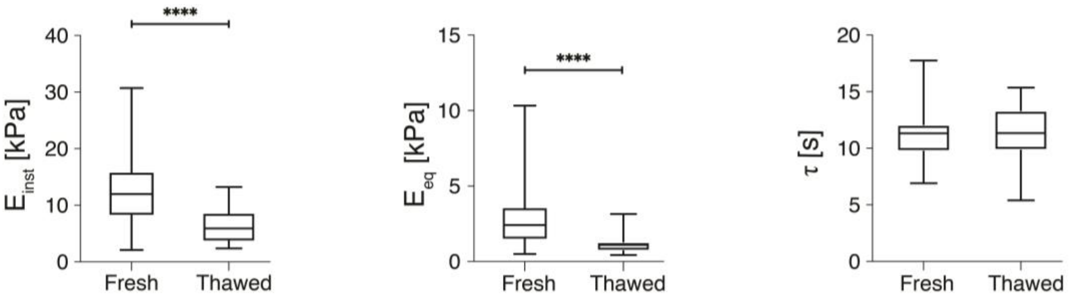


Fig. S1 Instantaneous elastic modulus ($E_{inst}$), equilibrium elastic modulus ($E_{eq}$), and characteristic relaxation time (τ) for all fresh and thawed samples tested with indentation stress-relaxation. **** = $p < 0.0001$

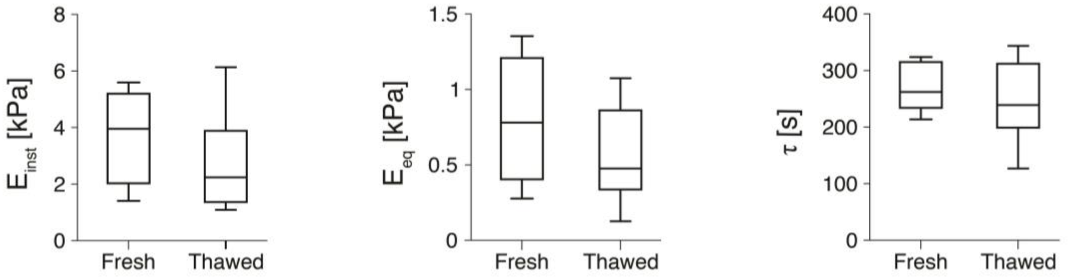


Fig. S2 Instantaneous elastic modulus ($E_{inst}$), equilibrium elastic modulus ($E_{eq}$), and characteristic relaxation time (τ) for all fresh and thawed samples tested with bulk stress-relaxation.

#### SI3. Excizing section influence – Indentation stress-relaxation

Since a significant difference was detected between fresh and thawed samples, the effect of cutting direction was investigated only on fresh samples, by means of all the typologies of mechanical tests mentioned in the main manuscript. As shown in Fig. S3, Fig. S4, Fig. S5, Fig. S6, Fig. S7, no statystical differences were reported for all the viscoelastic descriptors. Mann-Whitney was applied for each test, with the exception of bulk cyclic, for which Kruskal-Wallis was applied, followed by Dunn's post-hoc (GraphPad Prism 10). (*N* of samples per cutting section = 4; *N* of points per sample, for indentation = 15).

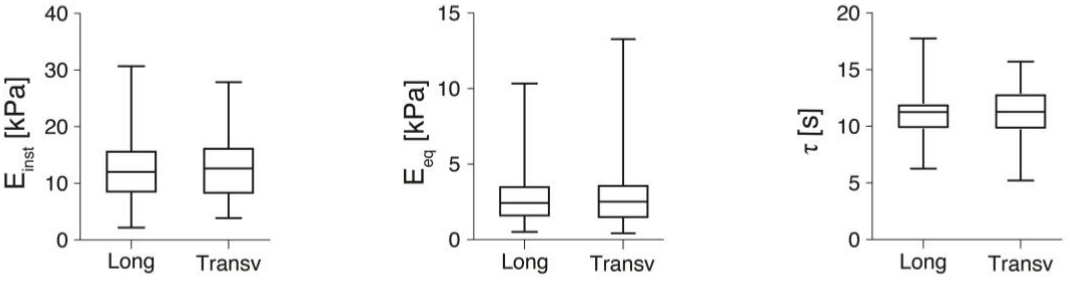


Fig. S3 Instantaneous elastic modulus ($E_{inst}$), equilibrium elastic modulus ($E_{eq}$), and characteristic relaxation time (τ) for all samples tested with indentation stress-relaxation and exctracted from the two considered cutting sections (longitudinal and transversal).

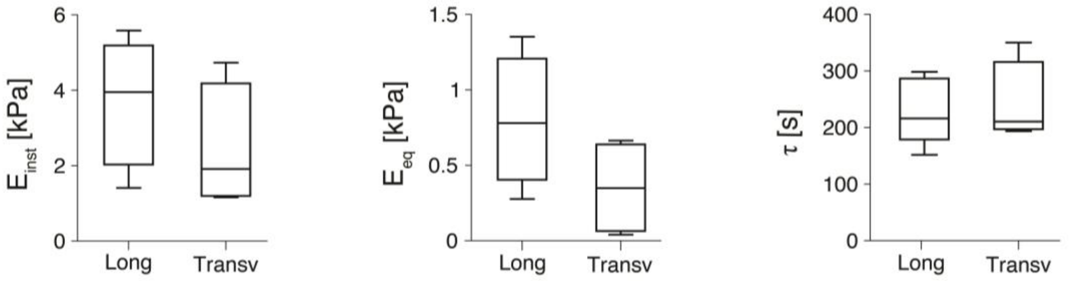


Fig. S4 Instantaneous elastic modulus ($E_{inst}$), equilibrium elastic modulus ($E_{eq}$), and characteristic relaxation time (τ) for all samples tested with bulk stress-relaxation and extracted from the two considered cutting sections (longitudinal and transversal).

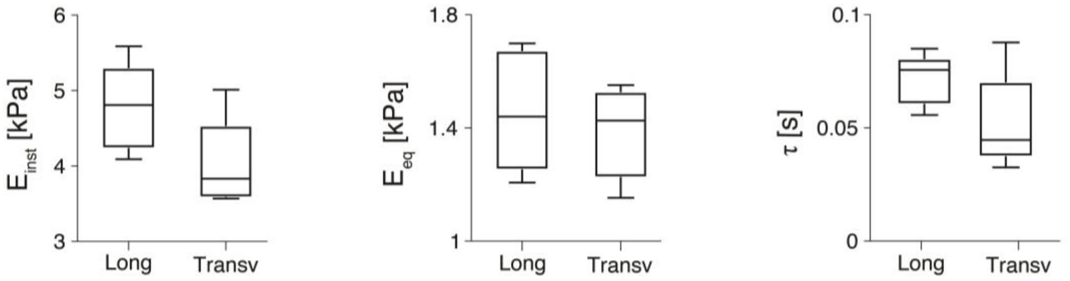


Fig. S5 Instantaneous elastic modulus ($E_{inst}$), equilibrium elastic modulus ($E_{eq}$), and characteristic relaxation time (τ) for all samples tested with bulk epsilon-dot spectrum and extracted from the two considered cutting sections (longitudinal and transversal).

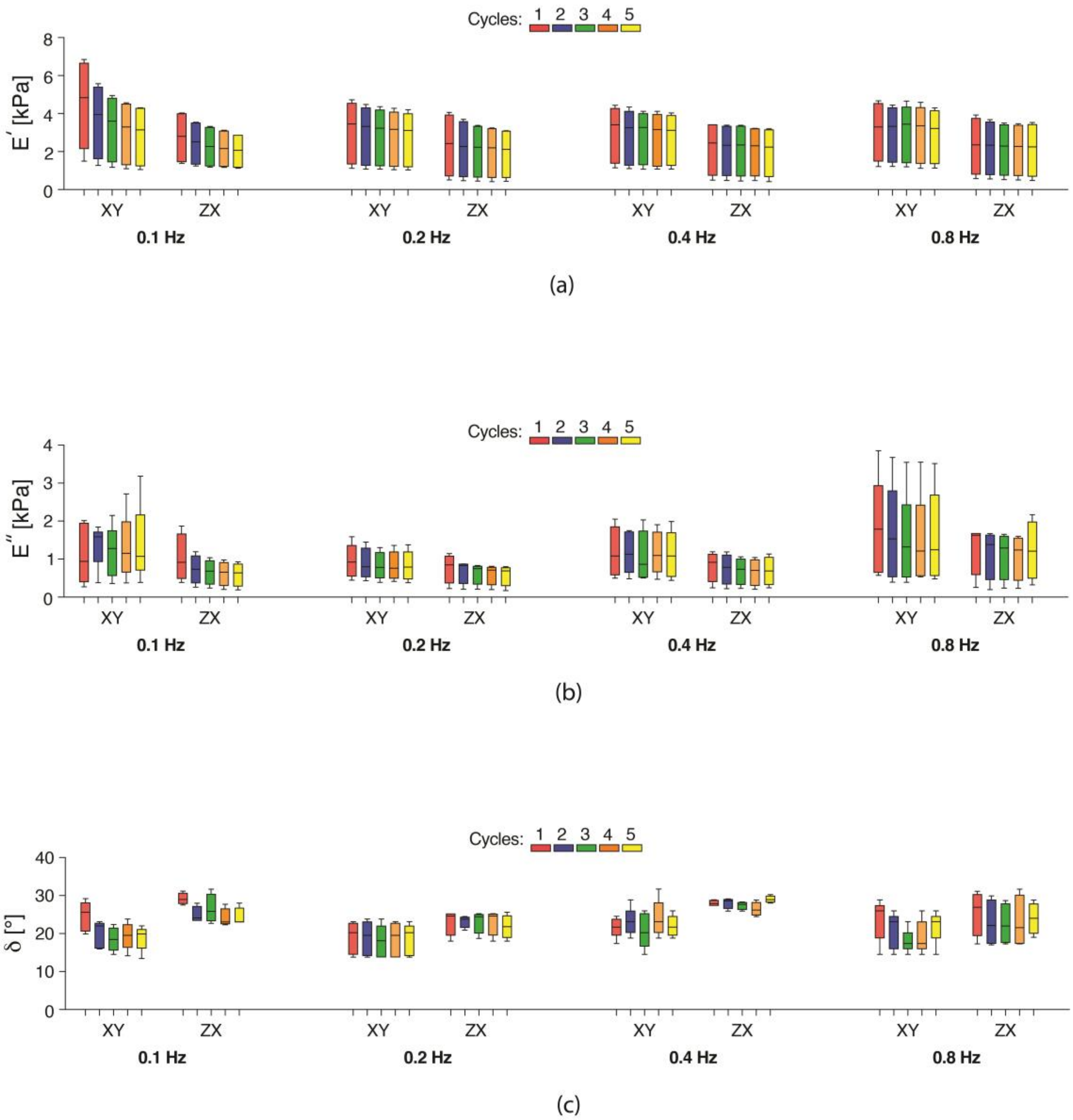


Fig. S6 Storage modulus (a), loss modulus (b) and phase shift (c) for all samples tested with bulk sinusoidal test (Frequencies: 0.1, 0.2, 0.4, 0.8 Hz; *N* of cycles = 5) and extracted from the two considered cutting sections (longitudinal and transversal).

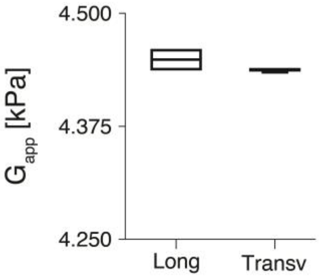


Fig. S7 Apparent shear modulus for all samples tested with shear and extracted from the two considered cutting sections (longitudinal and transversal).

##### SI4. Goodness of fitting with a second order ($n$=2) generalized Maxwell model – example from a bSR sample

| | $n = 1$ | $n = 2$ |
|---|---|---|
| **$E_0$ [Pa]** | 2556 | 2317 |
| **$E_1$ [Pa]** | 5333 | 3962 |
| **$\eta_1$ [Pa*s]** | 683437 | 853862 |
| **$E_2$ [Pa]** | - | 5946 |
| **$\eta_2$ [Pa*s]** | - | 57295 |
| **$R^2$ [-]** | 0.9177 | 0.992 |

##### SI5. Relaxation times for performed mechanical tests, expressed as median (25, 75%). i/bSR = indentation/bulk stress-relaxation; b$\dot{\varepsilon}$ = bulk epsilon-dot spectrum.

| | $\tau_1$ [s] | $\tau_2$ [s] |
|---|---|---|
| **iSR** | 0.3 (0.2, 0.4) | 11.2 (9.7, 12) |
| **bSR** | 7.3 (5.9, 9.9) | 215.5 (177.6, 287) |
| **b$\dot{\varepsilon}$** | 0.08 (0.06, 0.08) | x |

##### SI6. Shear and compression comparison

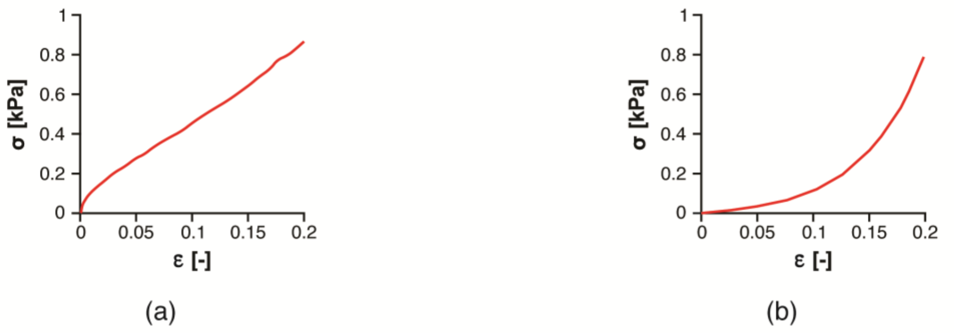


Fig. S8 Representative experimental curves for (a) bulk shear test and (b) bulk compression test. Both tests were conducted at the same flat indenter velocity (0.5 mm/s).

#### SI7. Representative experimental curves for each mechanical test

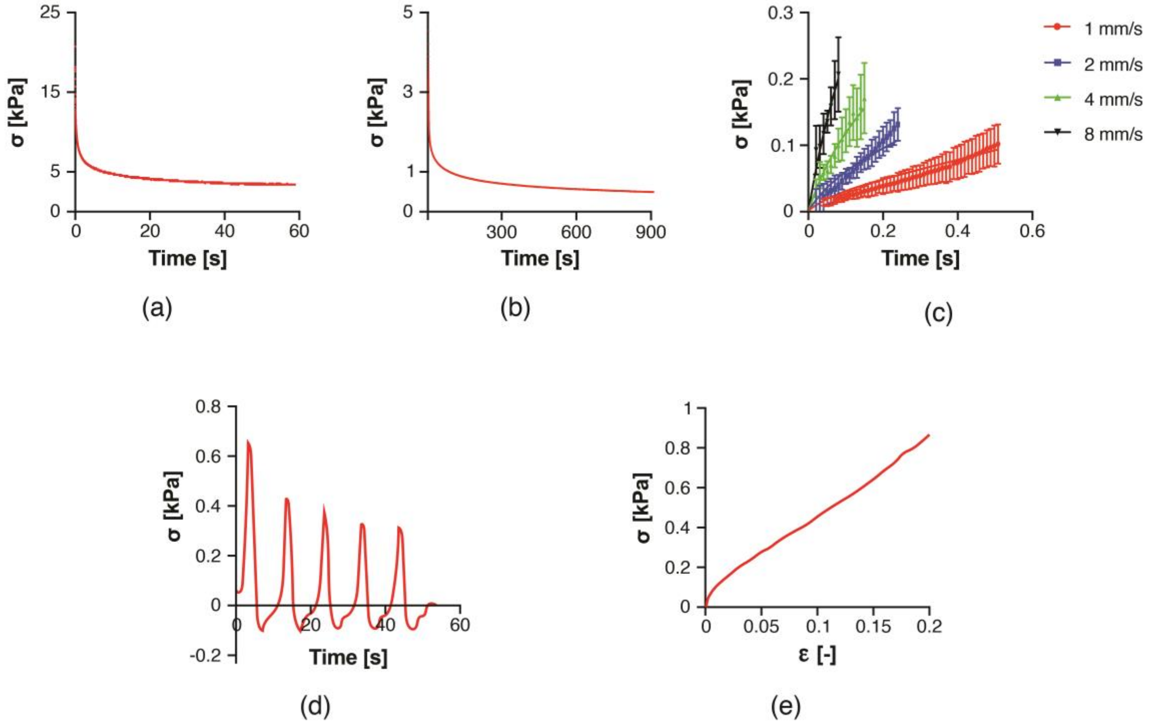


Fig. S9 Representative experimental curves for each performed mechanical test: (a) indentation stress-relaxation; (b) bulk stress-relaxation; (c) bulk epsilon-dot spectrum; (d) bulk sinusoidal; (e) bulk shear.
